# Stability of an adaptively controlled pathway maximising specific flux under varying conditions

**DOI:** 10.1101/550343

**Authors:** Gosse B. Overal, Josephus Hulshof, Robert Planqué

## Abstract

Microbial cells need to adapt to changing environmental conditions to survive. There is an evolutionary advantage to grow fast; this requires high metabolic rates, and an efficient allocation of enzymatic resources. Here we study a general control theory called *q*ORAC, developed previously, which allows cells to adaptively control their enzyme allocations to achieve maximal steady state flux. The control is robust to perturbations in the environment, but those perturbations themselves do not feature in the control. In this paper we focus on the archetypical pathway, the linear chain with reversible Michaelis-Menten kinetics, together with *q*ORAC control. First we assume that the metabolic pathway is in quasi-steady state with respect to enzyme synthesis. Then we show that the map between steady state metabolite and enzyme concentrations is a smooth bijection. Using this information, we finally show that the unique (and hence flux-maximising) steady state of this system is locally stable. We provide further evidence that it may in fact be globally stable.

## 1 Introduction

Microbes live in ever-changing environments to which they need to adapt in order to survive. If conditions are favourable, cells grow as fast as resources allow them to (Schaechter et al. 2006). Depending on the environmental conditions, cells use different metabolic networks to synthesise the components they are made of, using different sets of enzymes to catalyse the individual reactions in these networks. But even when the identity of the enzymes does not change, different resource availabilities, such as high or low concentrations of a food source, force cells to adapt the levels of the relevant enzymes. It is becoming increasingly clear that cells are indeed able to meet this challenge. They use enzyme resources economically (Basan et al. 2015, Bosdriesz et al. 2015, Li et al. 2014, Scott et al. 2014, You et al. 2013), and tune enzyme levels to maximise their growth rate (Dekel and Alon 2005, Jensen et al. 1995, Keren et al. 2016).

This adaptation to different environments is particularly surprising because many microbes do not have proteins in their membranes that would allow them to infer directly changes in resource concentrations outside the cell. Indeed, for microbes such as *Escherichia coli* and *Salmonella* that are able to grow on a multitude of carbon sources, having different membrane proteins to sense the presence of each resource would severely reduce the membrane area available for transport proteins. Instead, these microbes must rely on internal information about external changes. With changing external resource concentrations, internal metabolite concentrations must be used as proxies for those changes, for instance through metabolite-binding transcription factors influencing gene expression (Kochanowski et al. 2013, Kotte et al. 2010).

We have recently developed a general dynamical systems theory called *q*ORAC, or specific flux (*q*) Optimisation by Robust Adaptive Control that offers an implementation of this control problem (Planqué et al. 2018). It is formed by adding to a given metabolic pathway with prescribed enzyme kinetic rate laws a set of differential equations for enzyme synthesis. The details of the implementation are postponed to Section 2. The rates of enzyme production are constructed such that the only steady state of the combined metabolite-enzyme dynamical system is one in which the flux per unit expended enzyme, or “specific flux”, through the pathway is maximal. The optimal steady state flux attained depends on the resource concentration, but this concentration is not known in the enzyme synthesis control. Instead, the control uses an internal metabolite concentration as ‘sensor’. Some well-known example metabolites that act in this sensor role are Fructose-1,6-biphosphate (Kotte et al. 2014), allolactose (Gilbert and Müller-Hill 1966), and intracellular galactose (Sellick et al. 2008).

*q*ORAC-control may be added to any metabolic pathway with the property that it cannot be simplified. More precisely, this means that deleting any one reaction from the pathway would halt the flux through the pathway. Such metabolic networks are called Elementary Flux Modes (Schuster and Hilgetag 1994). The control is derived straight from the kinetic rate laws of the enzymatic reactions, and is designed to have the right steady state property of maximal steady state flux. There is no guarantee that solutions of the coupled dynamical system actually converge to this optimal state.

For each choice of external resource concentration the *q*ORAC-controlled linear chain has a unique steady state: the one in which steady state flux through the pathway is optimal (Planqué et al. 2018). Here we show, under a quasi-steady state (QSS) assumption that metabolic rates are much higher than enzyme synthesis rates, that this unique steady state is also locally stable, and provide additional insight that it might in fact be globally stable. Proving local stability requires us to first make an in-depth characterisation of the quasi-steady state, before focusing on the coupling with enzyme dynamics.

The structure of this paper is follows. First we review the *q*ORAC framework for general pathways—a detailed exposition may be found in (Planqué et al. 2018). Then we introduce the linear chain with reversible Michaelis-Menten reactions as the focal example. We proceed by assuming that metabolism is in quasi-steady state and study how metabolite concentrations in this QSS depend on the enzyme concentrations. Then we turn to the remaining slow enzyme dynamics and prove local stability. We finish with a discussion on global stability by considering a number of reasonable candidate Lyapunov functions, one of which is conjectured to be an actual Lyapunov function.

## 2 Maximising specific flux and the *q*ORAC framework

We start with the following model for the dynamics of metabolite concentrations. Let the vector ***x*** indicate all internal metabolite concentrations, 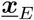 the vector of constant external metabolite concentrations, let ***v*** denote the vector of rates or fluxes of reactions in which metabolites are interconverted by catalyzing enzymes ***e***, and let the stoichiometric matrix be denoted by *N*. Then we consider as metabolic pathway the ODE system

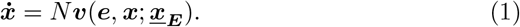

The functions 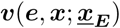 are assumed to be known and we make the assumption that each enzyme catalyzes exactly one reaction; in particular, we assume that 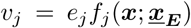. Such a linear dependence on enzyme concentration follows generally using QSSA-type analyses for enzyme-catalyzed reactions (Cornish-Bowden 2004).

Consider a choice of enzyme concentrations ***e*** with total enzyme concentration 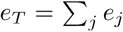 such that it allows a steady state flux through the pathway. Then, by scaling all enzyme concentrations by the same constant, a new steady state with the same metabolite concentrations may be constructed, with a flux exactly scaling with the same factor. In other words, the flux per unit of total enzyme concentration, or specific flux, remains constant. The linear dependence of reaction rates on enzyme concentrations also implies that maximising the steady state flux for a given total enzyme concentration is equivalent to minimising the total enzyme concentration necessary to attain one unit of steady state flux. (To clarify, the analogy in which trying to buy the maximal number of apples for 10 EUR is equivalent to trying to buy the cheapest apples is here an *exact* analogy.) For the purposes of this paper, it turns out that it is more fruitful to consider the question of maximal flux per unit total enzyme than it is to minimize enzyme levels for one unit of flux.

It has recently been shown (Müller et al. 2014, Wortel et al. 2014) that maximal steady state specific flux is attained in special type of pathway, called an Elementary Flux Mode (EFM). An EFM is a pathway that has a minimal number of enzymes involved: it allows for a balanced flow of metabolism, but this possibility is void if any one of its enzymes is removed from the pathway. As a result, if the pathway is in steady state and one flux is known, then all fluxes are determined. An EFM may hence be denoted by a fixed vectors ***C***, in which one flux value, for instance the target flux that is to be maximised, is set to 1. Any other steady state flux through the EFM may be characterised by ***v*** = *c**C***, with *c* the target flux. A more in-depth description of EFMs may be found in (Papin et al. 2004, Schuster and Hilgetag 1994, Schuster et al. 2002).

A given EFM, however, still allows many steady states: any choice of positive enzyme concentrations that take part in the EFM generally gives rise to a corresponding vector of steady state metabolite concentrations, and the resulting steady state flux is not maximal unless the enzyme allocation is chosen exactly right.

Consider an EFM with *n* reactions, each with steady state rate *v*_*i*_ = *cC*_*i*_ = 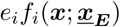. Then

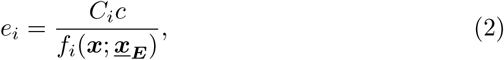

where *i* = 1, …, *n* indexes the *n* enzymes, and

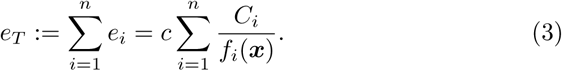

So for a fixed *e*_*T*_, the steady state flux *c* is a function of only the metabolite concentrations

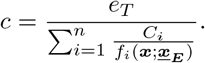

Thus finding the maximum *c* can be reformulated as finding the vector ***x***, given the external conditions 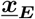, that minimises the objective function

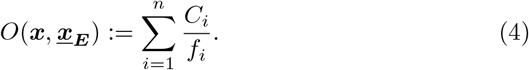

It was recently shown that for fixed external concentrations 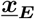, the objective function (4) has a unique minimum ***x***^*o*^ for a large class of rate laws (Noor et al. 2016, Planqué et al. 2018).

Any minimiser ***x***^*o*^ of 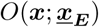 is also the unique critical point of this objective function, and hence solves the *optimum equations*

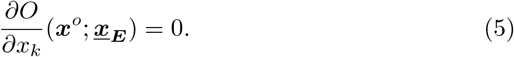

Rather than prescribing 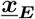 in (5) and calculating the remaining variables ***x***^*o*^, one might also prescribe some internal variables, termed sensors, ***x***_*S*_, and solve for all other variables (including the external concentrations). The resulting object is called the *optimum as predicted by the sensors*, or predicted optimum in short. The Implicit Function Theorem gives the requirements which internal metabolites may be used for this procedure. Locally, the sensors should allow a parametrisation of the family of optima that was initially parametrised by 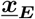. In particular, 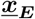 and ***x***_*S*_ must have the same number of elements. The optimum predicted by ***x***_*S*_ is denoted by ***ξ***(***x***_*S*_). Clearly, if 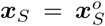, then 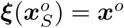.

The estimated optimal enzyme distribution ***ε*** can then be computed from ***ξ*** through the steady state equations of metabolism (2). Denoting the kinetic functions *f*_*j*_ as

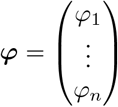

if they use ***ξ*** as argument rather than ***x***, then

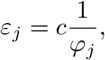

where *c* is such that the total amount of enzyme (3) is *e*_*T*_. Without loss of generality, we assume *e*_*T*_ = 1, by setting

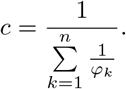

In conclusion, based on the sensor concentration ***x***_*S*_, the estimated optimal enzyme distribution is

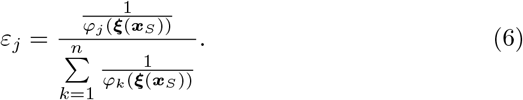

We can now supply to the dynamical system for the metabolic pathway (1) a set of differential equations for enzyme concentrations involved in the pathway. The general structure of such equations is assumed to be

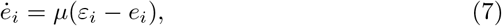

where *µε*_*i*_ describes the enzyme synthesis rate of enzyme *i*, and the degradation term involves dilution by growth in a cell population growing at rate *µ*. (This last term could have been present in (1) as well, but is neglected because *µ* ≪ *v*_*j*_ in biological systems.)

To summarise, the complete dynamical system for a *q*ORAC-controlled pathway is given by

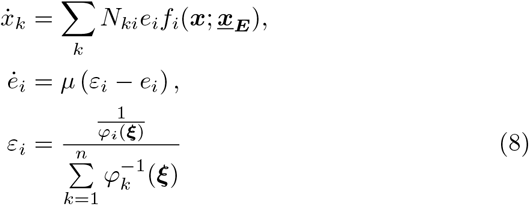

and ***ξ*** is defined by

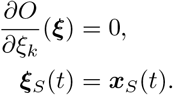

The construction ensures that if this system converges to a steady state, it is necessarily one with maximal steady state flux (Planqué et al. 2018). Since the enzyme synthesis rates do not depend on 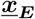, this pathway is robust to changes in 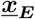. The reason is essentially that the complete dynamical system can only be in steady state if the sensor has the right concentration, and therefore predicts the right optimal enzyme concentration.

It is of course far from clear that this dynamical system in fact does converge to steady state. In this paper we show local stability of the steady state for the most important and also simplest EFM, the linear chain of reversible reactions, under the additional assumption that metabolism is at quasi-steady state.

### 2.1 *q*ORAC for the linear chain

The analysis of this paper is confined to the study of the archetypical EFM, the linear chain with *n* enzymatic reactions,

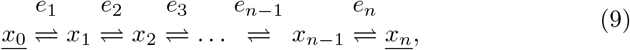

where external nutrient 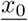 is converted into the the external waste 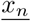. The stoichiometric matrix is given by

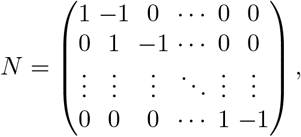

which has a one-dimensional null space spanned by the vector

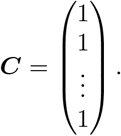

The rate functions are assumed to be given by standard Michaelis-Menten kinetics, in which we choose kinetic constants as follows,

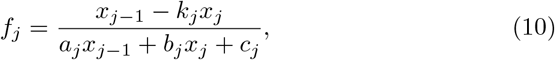

where for *f*_1_ and *f*_*n*_ we have 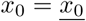 and 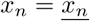.

For the remainder of this paper we assume that 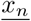 is a fixed parameter and that the external nutrient concentration 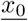 may vary.

#### Definition 1

*The nutrient concentration 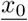 and waste concentration 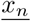 are* positively oriented *if a positive steady state flux through the linear chain is possible, which follows exactly if*

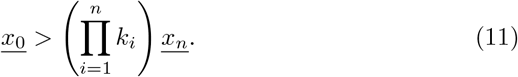

Every positively oriented 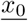 yields a unique minimum ***x*** of 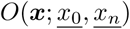. We denote the set of all minima for all positively oriented 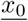 as

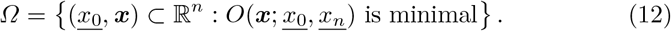

We now show that in principle, any internal metabolite could be used as a sensor in the *q*ORAC-control. In other words, for the linear chain *Ω* can be parametrised by any internal metabolite.

#### Lemma 2

*Let x*_*s*_ *be positively oriented with respect to 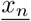 for some* 1 ≤ *s* ≤ *n* − 1. *Then there is a unique*

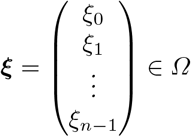

such that *ξ*_*s*_ = *x*_*s*_

*Proof* Since *x*_*s*_ is positively oriented, 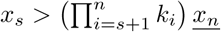. Metabolite concentrations *x*_*s*_ and 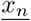 are the positively oriented endpoints of a linear (sub)chain with enzymes *e*_*s*+1_, …, *e*_*n*_, and therefore the function

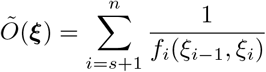

has a unique minimum for *f*_*i*_ > 0. This minimum has *ξ*_*s*_ = *x*_*s*_ and is the unique solution to

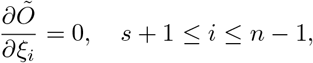

where *f*_*i*_ > 0. For *Ω*, the defining equations are

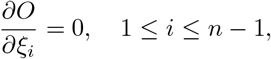

where *f*_*i*_ > 0. The first *s* functions *f*_1_, …, *f*_*s*_ depend only on *x*_1_, …, *x*_*s*_, so the minumum of *Õ* is the unique solution to the subset of defining equations for *Ω*,

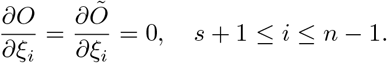

Hence, any solution ***ξ*** Ω *Ω* with *ξ*_*s*_ = *x*_*s*_ has these values for *ξ*_*s*_, …, *ξ*_*n*_−_1_.

Note that

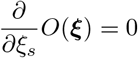

is an equation in *ξ*_*s*−1_, *ξ*_*s*_ and *ξ*_*s*+1_ only, with *ξ*_*s*_ and *ξ*_*s*+1_ already known. With the prescribed kinetics (10), the resulting equation for *ξ*_*s* − 1_ is a quadratic polynomial. It has two solutions, exactly one of which has *f*_*s*_(*ξ*_*s*−1_, *ξ*_*s*_) > 0.

By determining the subsequent coordinates in sequence, each time taking the larger of the two solutions of the polynomial we need to solve, we construct a solution to the equations

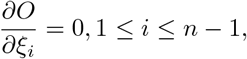

where *f*_*i*_ > 0 ensures that there is always but one choice. Therefore this solution is unique.

The implementation of (8) for the linear chain is given by,

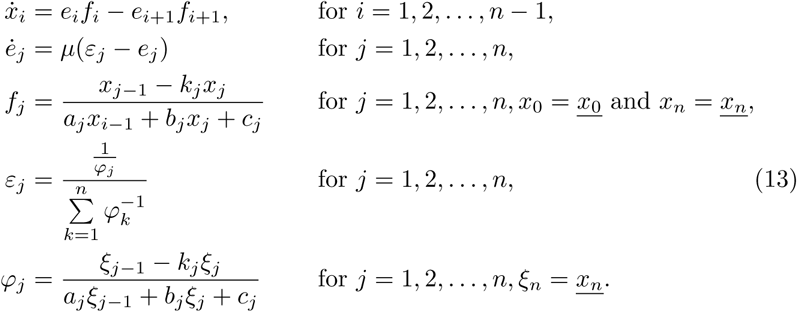

The sensor concentration *x*_*s*_(*t*) defines the estimated optimal steady state metabolic concentrations ***ξ***(*t*) as the unique element ***ξ** ∈ Ω* that satisfies *ξ*_*s*_ = *x*_*s*_(*t*). Note that this includes the external nutrient 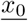 that is sensed for, estimated as *ξ*_0_.

To aid the reader, all relevant vectors are explicitly given below,

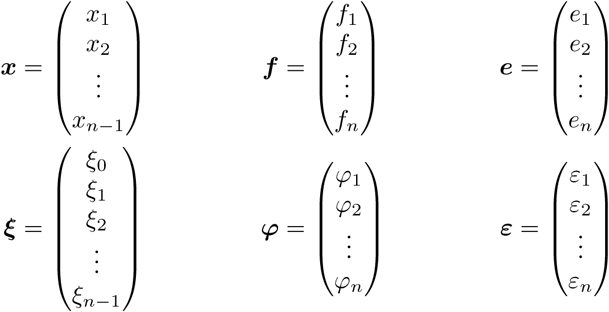

## 3 Results

### 3.1 Timescale separation

We assume that the metabolism rates are much higher than the enzyme production and dilution by growth rates, i.e., *µ* is a small parameter. Hence we separate the timescales.

#### 3.1.1 Fast timescale

For the fast timescale we set *µ* = 0 and thus consider the enzyme concentrations *e*_*i*_ to be static while the metabolism flows. The differential equations are then given by

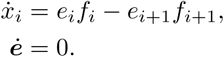

From (Smillie 1984) we know that the linear chain without enzyme dynamics has a unique steady state that is globally stable. Since this older result was not known to us at the start of our investigation, we first gave a different proof of global stability ourselves, which is supplied in the Appendix.

Thus in this timescale, the metabolite concentrations ***x*** will converge to a unique solution, 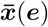.

#### 3.1.2 Slow timescale

The slow timescale follows from substituting *τ* = *µt* in the original system. Then the time derivative changes as

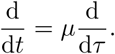

The differential equation system changes to,

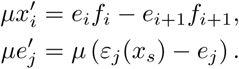

Dividing out *µ*, we have

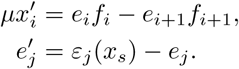

Setting *µ* = 0, we get the differential algebraic system that defines the dynamics of the slow timescale for the quasi equilibrium 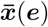 and the time-dependent enzyme concentrations. They are

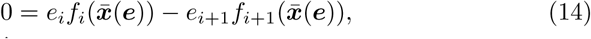

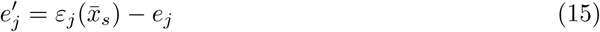

for *i* = 1, …, *n* − 1.

#### 3.1.3 Explicit dependence of the metabolic Quasi Steady State on enzyme concentrations

We rewrite these equations to a form more amenable to analysis, by adding the steady state flux as an extra variable *c* as follows. For any ***e***, the solution 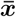 yields that

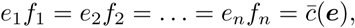

where the extra equation *e*_*n*_*f*_*n*_ = *c* adds a dependent variable *c*, from which we can rewrite all the other steady state equations to

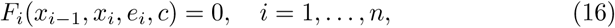

where

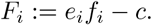

Now we have *n* equations in *n* variables (***x*** and *c*). For any ***e***, the functions 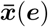 and 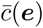 are such that equations (16) are equivalent to (14) and solve

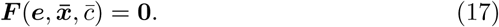

These alternative equations yield a clearer picture of how 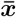 depends on ***e***, which we will deduce step by step. To be more precise, we will first introduce a partial solution ***x***^∗^(***e**, c*) based on *n* − 1 equations (*F*_2_ = 0*, …, F_n_* = 0) and derive explicitly its partial derivatives to ***e*** and *c* (Lemma 3). Then we solve the last equation (*F*_1_ = 0) for *c*, yielding 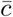. We find explicit derivatives of 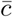 to ***e*** (Lemma 4). Combining this with the results for *x*^∗^, we calculate partial derivatives of 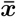 to ***e***.

For convenience, we introduce some notation for the partial derivatives of the flux functions to their substrate and product concentrations,

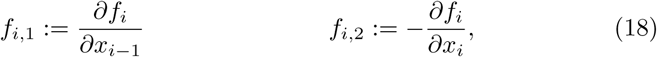

where *i* = 1, …, *n*. Note that *f*_*i*,1_ > 0 and *f*_*i*,2_ > 0, but that *f*_1,1_ and *f*_*n*,2_ should be disregarded, because 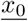 and 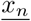 are not dynamic variables. Anywhere where we do write *f*_1,1_, it will be both in the numerator and denominator of a fraction rendering it irrelevant; this is done only to make the notation uniform.

We can immediately see which terms are positive and negative in the partial derivatives of *F*_*i*_ = *e*_*i*_*f*_*i*_ − *c*:

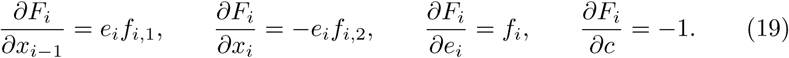

In the derivations to come we often come across the following terms,

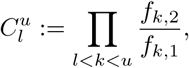

where *l* ≤ *u* − 2. As we can see that

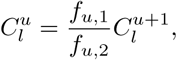

we can generalise this notation also for when *l* ≥ *u* − 1,

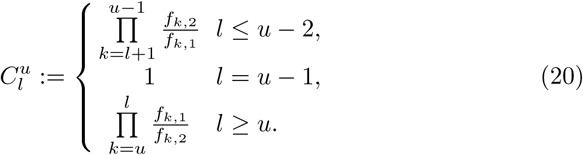

This will make the expressions for the explicit partial derivatives of ***x***^∗^, 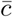 and 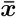 more convenient.

##### Lemma 3

*For any **e*** > **0** *such that* 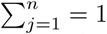 *and c* > 0 *small enough, there exist unique solutions* 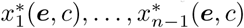, *that solve F*_2_ = 0, …, *F*_*n*_ = 0. *Furthermore their partial derivatives are:*

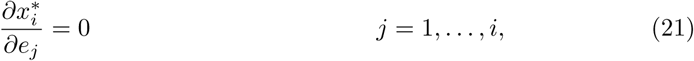

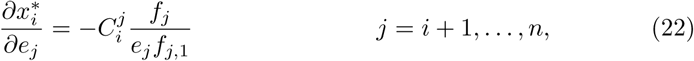

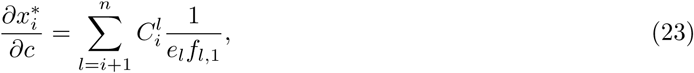

*for i* = 1, …, *n* − 1, *with* 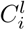 *and* 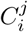 *given by* (20).

*Proof* A sketch of the argument that this proof is based on can be found in Figures 1 and 2.

**Fig. 1:**
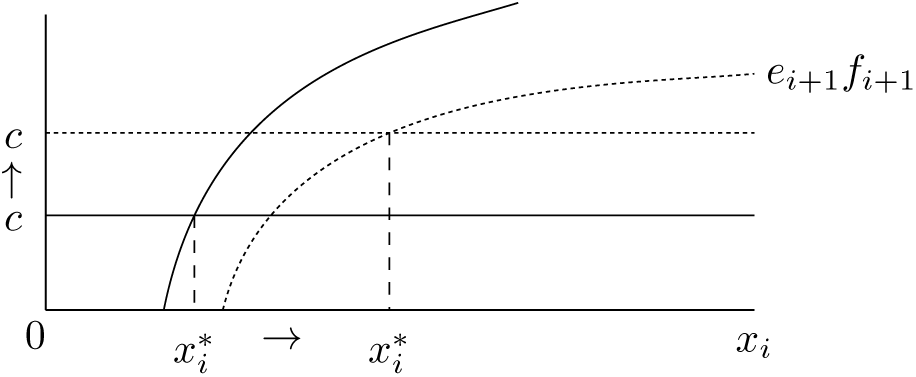
Schematic representation of F_*i*+1_ = 0 and how the solution 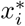 depends on c. For increased c (the dotted graphs), f_*i*+1_ decreases, the equilibrium value 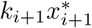 is increased, thus the solution 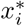 is increased.

**Fig. 2:**
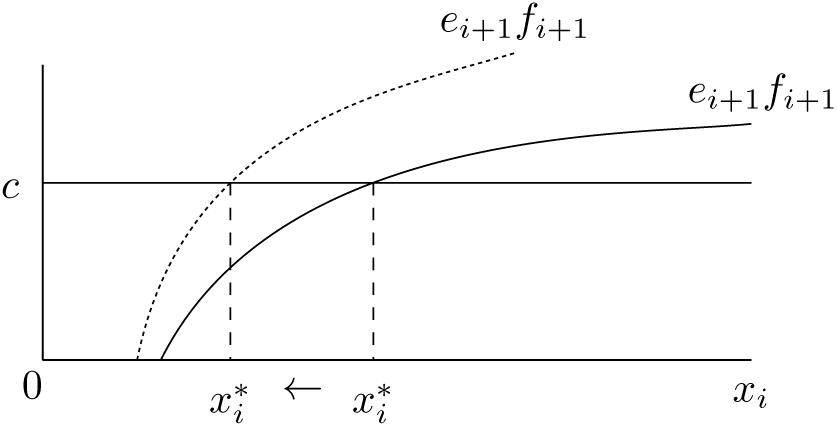
Schematic representation of F_*i*+1_ = 0 and how the solution 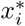 depends on e_*j*_ for *j* ≥ *i* + 1. For increased e_*j*_ (the dotted graph), 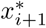 decreases, increasing f_*i*+1_ and the solution 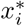 becomes smaller. Note how the dotted graph is not just above the original graph, but that the chemical equilibrium value 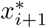 where f_*i*+1_ = 0 is decreased due to e_*j*_ changing the base value 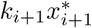. If *j* = *i* + 1, however, the dotted graph should actually have the same origin on the horizontal axis, and an increased e_*i*+1_ would still put the dotted graph above the original.

Let 1 ≤ *i* ≤ *n* − 1. If *i* = *n* 1, the next metabolite is external, 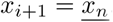. Otherwise assume that 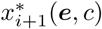 solves *F*_*i*+2_, such that

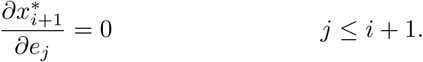

Considering *F*_*i*+1_, note that if *x*_*i*_ is at the boundary of the admissible domain, 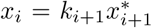, then *f*_*i*+1_ = 0 and *F*_*i*+1_ = −*c*. On the other end, *x*_*i*_ is not bounded and *f*_*i*+1_ saturates to some maximum value. If *c* is smaller than the maximum vlue of *e*_*i*+1_*f*_*i*+1_, *F*_*i*+1_ > 0 follows for great enough *x*_*i*_. Furthermore,

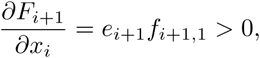

so inbetween these extremes, there is a unique 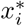 that solves 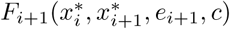, by continuity.

We consider the dependency of *F*_*i*+1_ on its variables,

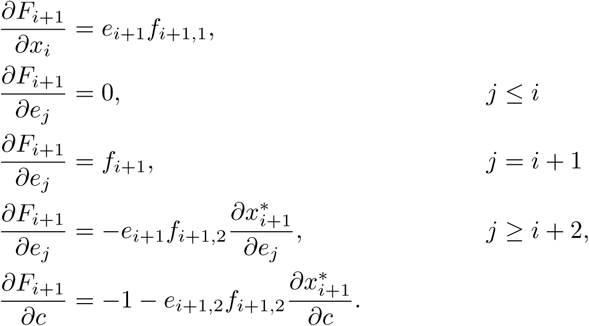

Note the difference with (19) that comes from using the implicit solution 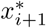 with the chain rule. Partial differentiation then yields the recursively defined derivatives by the Implicit Function Theorem, assuming that *c* is small enough.

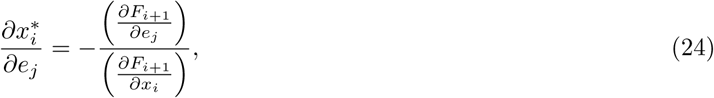

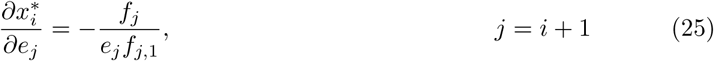

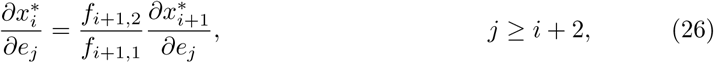

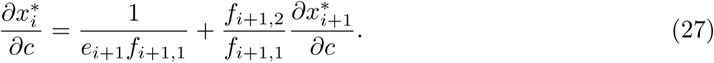

For any pair *i* = 1, …, *n* − 1, *j* = 1, …, *n*, either 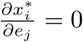 or solution (26) can be iteratively applied, until *j* = *i* + 1 (solution (25)), yielding the explicit partial derivatives

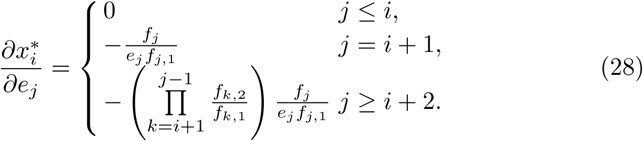

For any 1 ≤ *i* ≤ *n* − 1, solution (27) for the *c*-dependence of 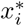 can also be iteratively applied, taking into account that 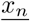 is constant, so

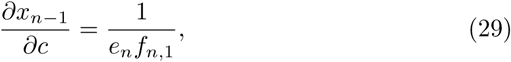

yielding the explicit partial derivatives

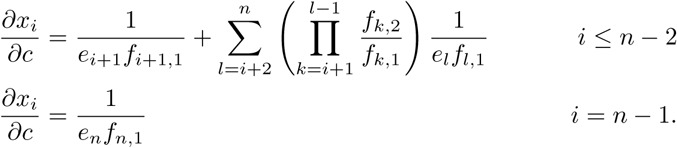

If we now substitute (20) into the equations, the derivatives are given by (21), (22) and (23).

Note that the 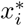 do not depend on *e*_1_ as it always falls under (21), because 1 ≤ *i* for all *i*.

We are ready to solve the last equation *F*_1_ = 0 for *c*, using 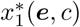,

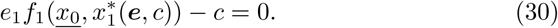

##### Lemma 4

*For any **e*** > **0** *such that* 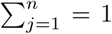, *there is a unique* 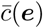 *that solves* (30), *such that together with* **x**^∗^ *it represents the slow manifold*, 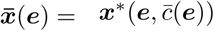. Furthermore the partial derivatives of 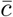 are given by

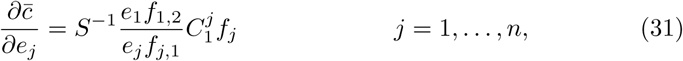

*where* 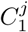 *is given by* (20) *and S*^−1^ *normalises the factors in front of the f*_*j*_ in the partial derivatives above,

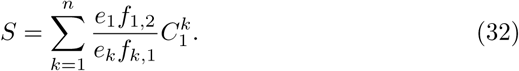

*Proof* Equation (30) has the following partial derivatives, based on the known partial derivatives of 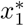 (21), (22) and (23),

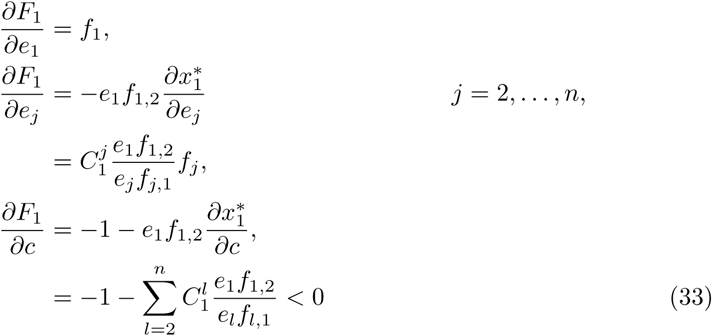

If *c* = 0, the flux functions are at chemical equilibrium, 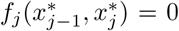, so 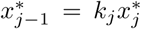, for *j* = 2, …, *n*, given that ***e*** > 0. Therefore the intermediate solution is then at equilibrium with the waste concentration 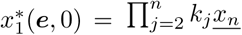 and

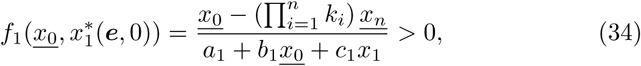

because we have assumed our system to be positively oriented (11).

This proves that if *c* = 0, 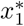 is such that *F*_1_ > 0.

On the other end of the spectrum, 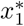 is only defined for *c* small enough and as *c* approaches this bound, 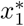 will become unbounded. In particular for *c* close to this bound, we get

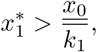

so *F*_1_ is negative for *c* large enough.

Note that *F*_1_ is decreasing in *c* (33) and goes from positive to negative over the domain of *c*. Therefore there is a unique solution 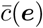 by continuity. Substituting 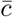 into ***x***^∗^ yields a unique solution to the steady state equations that define the slow manifold 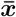 (17).

Furthermore from the Implicit Function Theorem, we get the partial derivatives are given through implicit differentiation using the above equations and Lemma 3,

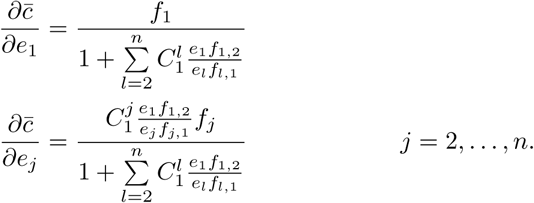

If we use a mathematical trick to substitute

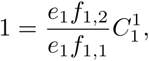

the partial derivatives are given by

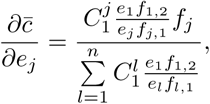

for *j* = 1, …, *n*, which is exactly what we wanted to get (31) if we substitute *S* (32). This trick is why 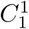 was defined as it was (20) and why *f*_1,1_ can still be disregarded.

To get some intuition for the flux *c* in the equations of the argument above, consider some intermediate enzyme with concentration *e*_*j*_. It is only involved in the equation *f*_*j*_. If we set *c* = 0 in this equation, we force the flux through this enzyme to zero, yet if we push *c* to its maximum value that still has a solution in *f*_*j*_, the substrate concentration is pushed up to infinity. This is in particular beyond its maximal value where it is at chemical equilibrium with the nutrient concentration 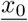. That forces all enzymatic fluxes leading up to *e*_*j*_ to be zero, so *c* is caught inbetween being zero in this equation and pushing itself to zero in other equations by increasing in this equation. The Lemma above shows that if *e*_*j*_ > 0 for all *j*, we can push *c* up from zero in all instances at once to then push 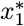 from chemical equilibrium with 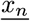 up to 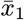, leading to the steady state flux 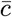.

##### Corollary 5

*The implicit function* 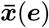 *that defines the slow manifold has the following partial derivatives*,

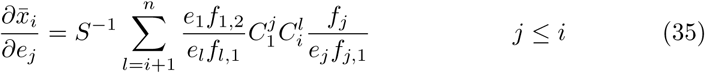

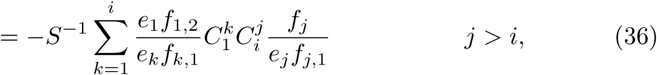

*where S is given in* (32)

*Proof* The function 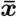 follows from ***x***^∗^ and 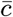,

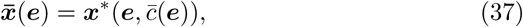

thus for any *i* = 1, …, *n* − 1 and *j* = 1 …, *n* it follows that

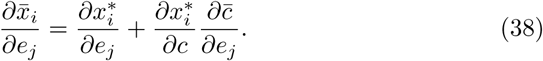

Taking the expressions from the derivatives given in (23) and (31), we can immediately see

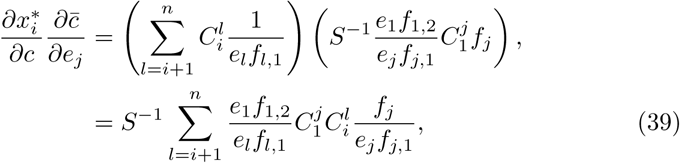

with *S* as in (32), in which 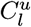 is defined in (20).

From (21) and (22), we recall

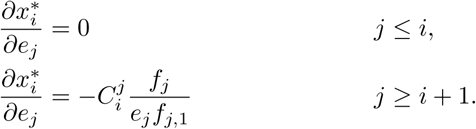

Hence for 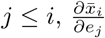 is given by (39).

For *j ≥ i* + 1 we have to do some more work, but we recognise in (39) we can rewrite the following

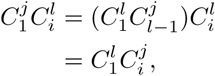

for *l ≤ j*. Otherwise the same identity holds,

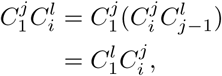

for *l ≥ j* + 1, manipulating the factors based on the definition (20).

Combining these expressions in (38) and recalling *S* from (32) yields

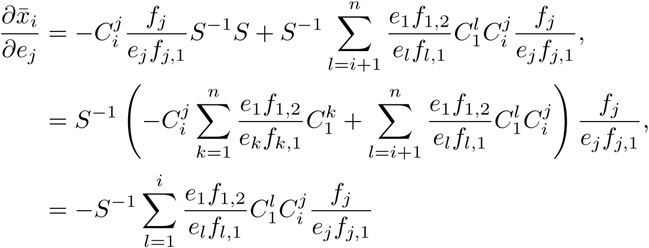

From the explicit form of the partial derivatives of 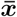 to all ***e***, we can see what is the time-dependency for 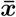.

**Corollary 6** *In the slow timescale, we have*

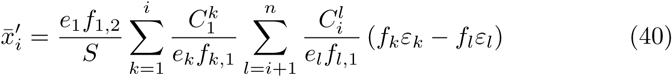

*Proof* The change of an internal metabolite concentration in the slow timescale completely depends on the change in the enzyme concentration,

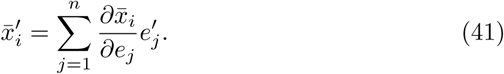

The expression for the partial derivative 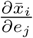 is qualitatively different for *j ≤ i* (35) and *j > i* (36). Hence we split the sum (41) into these two parts, *j*_1_ ≤ *i* and *j*_2_ > *i*.

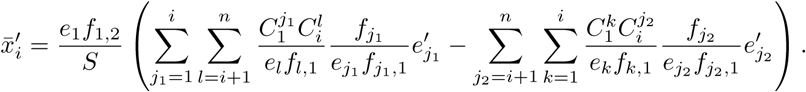

We can rename the indices *j*_1_ = *k*, *j*_2_ = *l* and see that the two double sums can be integrated,

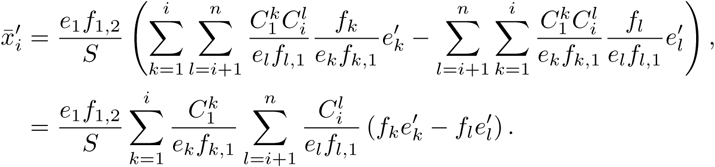

The expressions for the time derivatives of the enzyme concentrations (15) can be substitited and as we are at QSS, the steady state equations hold,

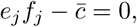

for *j* = 1, …, *n*, therefore we can finish the proof to the desired expression,

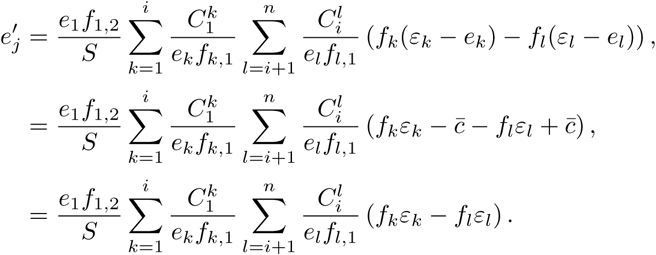

From Lemma 4 and Corollary 5, we get derivatives towards the direction of increasing *e*_*j*_, which does not comply with the notion that the total amount of enzyme remains constant at *e*_*T*_ = 1. To interpret the partial derivatives inside the space where *e*_*T*_ = 1 holds, we would have to make some linear combination of the *e*_*j*_ dependent on the other variables and introduce directional derivatives. A simple way is to take 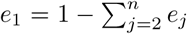 for instance, such that increasing any enzyme concentration other than *e*_1_ means decreasing *e*_1_ by that much and conserving the total amount.

One key observation is that in the direction

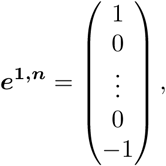

the total amount of enzyme is conserved and the directional derivative is positive

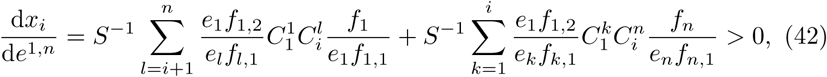

for all *i* = 1, …, *n* 1. This has the consequence that the subspace of the enzyme distribution space defined by having some fixed *x*_*i*_ is a smooth manifold.

### 3.2 The Quasi Steady State defines a smooth bijection

So far we have assumed that the enzymes are all present, *e*_*j*_ > 0 for all *j*, which is not unreasonable, but there is some insight to be gained from considering the boundary where some enzyme concentrations are zero.

If exactly one enzyme concentration is zero, *e*_*j*_ = 0, the function 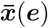 is also well-defined. Obviously then there is no flux through the system: *c* = *e*_*j*_*f*_*j*_ = 0, and because all other enzyme concentrations are positive, *e*_*k*_ > 0, *k* ≠ *j*, the remaining fluxes must be at chemical equilibrium, *f*_*k*_ = 0, *k* ≠ *j*. Thus the first few metabolite concentrations (*i* ≤ *j* − 1) are at equilibrium with the nutrient concentration 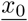 and the last few metabolite concentrations (*i ≥ j*) are at equilibrium with the waste concentration 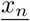. Therefore 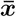 is uniquely defined for these border-points, but it is not an injection, as the exact values of ***x*** are now given, solely based on the information that exactly one enzyme concentration is zero, *e*_*j*_ = 0 < *e*_*k*_, for some *j* and all *k ≠ j*.

Let us therefore define clearly the spaces for which we will show that 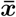 is a bijection,

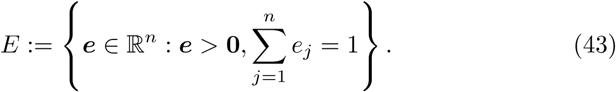

The function 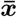, implicitly defined solving equations (17), is a bijection between *E* and the space of admissible metabolite concentrations

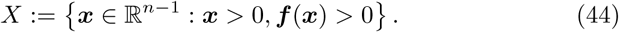

#### Lemma 7

*For the nonzero enzyme concentrations, the steady state metabolite concentrations are uniquely defined and they uniquely define the enzyme profile. The implicit function*

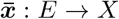

*defined by equations* (17) *is a smooth bijection.*

*Proof* Note that from Lemma 4, this function is well-defined for all *e ∈ E*. We have already shown in (6) that it has a unique inverse, given by

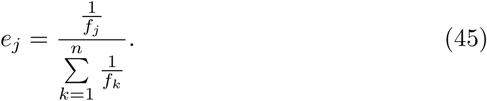

To prove smoothness, note that we have shown that inside *E* there is always a direction in which all *x*_*i*_ increase (42), hence the Jacobian matrix of 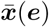 always has full rank. Smoothness follows from the Implicit Function Theorem.

### 3.3 Local stability

In the Appendix we show that it is possible to rewrite the dynamical system to one for the metabolites only, see Theorems 12 and 13. However, for the consideration of local stability it makes more sense to avoid this ‘simplification’, because the direction of flow in the space of enzymes is much simpler,

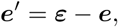

even though the definition of 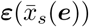 is not simple at all. Nevertheless, as we will see, detailed knowledge of the quasi-steady states will become informative in the proof of local stability.

There is a subtle and important difference between the steady state for (***ξ**, **ε***) and for 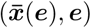. In the case of ***ξ***, the external concentration *x*_0_ is estimated at *ξ*_0_(*x*_*s*_), and ***ξ*** minimises the objective function *O*(***x***; *ξ*_0_); in the case of 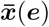, the nutrient concentration is fixed at its actual value 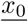, while it does not necessarily minimise the objective function. As the estimated optimal enzyme profile ***ε*** has sum equal to 1 by definition (13), it follows that ***ε** ∈ E*.

One could consider 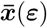 and compare this to ***ξ***. If ***ε*** and *ξ*_0_ are known, we can retrieve ***ξ*** through the steady state equations. However, due to the subtle difference between 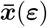 and ***ξ***, we have to retrieve it with a shifted external nutrient concentration *x*_0_ = *ξ*_0_.

The equations governing these two maps from ***ε*** are so similar that we can predict an ordering between the two, based solely on how accurate the sensor concentration *x*_*s*_ is. If the current sensor value is different than the sensor in optimum, the sensor value 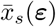 at the predicted enzyme distribution ***ε*** is in the same direction as the optimum.

Let 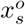 denote the sensor concentration in the unique optimum (recall *Ω* (12)) as defined by 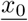.

#### Lemma 8

*For any **e** ∈ E*,

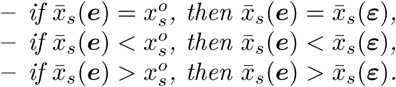

*Proof* Let ***e** ∈ E*. The corresponding point on the slow manifold 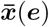 has sensor value 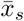, which yields ***ε***.

If 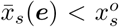, then ***ξ*** = ***ξ**^o^*, 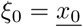 and ***ε*** is the true optimum. Therefore

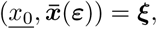

so it follows that

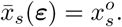

This proves the first result of the lemma.

Next, we assume 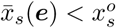. The set *Ω* defined in (12) has the property that if *x*_*s*_ increases, the corresponding element ***ξ** ∈ Ω* increases in all of its entries. This is a corollary to Lemma 2 as any alternative to this would contradict that result. Specifically this means that *ξ*_0_ increases with *x*_*s*_ and from 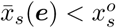 it follows that 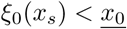.

The intermediate solution ***x***^∗^(***ε**, c*) yields ***ξ*** from ***ε*** if the following equation holds for *c*,

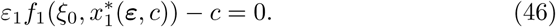

We recall that for 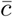 equation (30) holds,

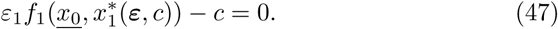

If (47) holds, the intermediate solution yields 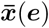. Define c = 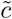 as the solution to (46). Then the intermediate solution yields ***ξ***. Note that the only difference between (46) and (47) is that we consider *x*_0_ = *ξ*_0_ instead of *x*_0_ = 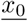 respectively.

So both 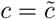 and 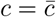 are solutions to equation

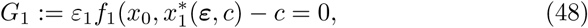

for *x*_0_ = *ξ*_0_ and 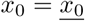 respectively.

Partial differentiation of (48) yields that

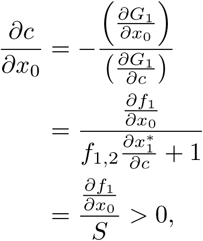

where we use the known expression for 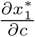 (23). So 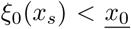 yields that 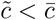.

Note that the statement that 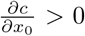 follows immediately from its interpretation as well: if the nutrient concentration increases, the steady state flux of balanced metabolism increases.

Recall that 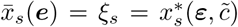 and 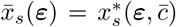. From expression (23) in Lemma 3, it follows that the intermediate solutions increase with *c*, 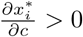, *i* = 1, …, *n* − 1. Specifically the intermediate solution for the sensor increases with *c*, 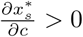, we conclude that 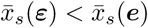.

For 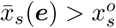 the proof follows in the same way.

This ordering is rather abstract, but strong in consequence. Combining it with the result of Lemma 7 that 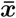 defines a smooth bijection, this proves that the unique and optimal steady state of the linear chain is locally stable.

#### Theorem 9

*The unique steady state **ε***^*o*^ *of qORAC for the linear chain of n enzymes is locally stable in the slow timescale*.

*Proof* The optimum ***ε***^*o*^ is the given from ***ξ***^*o*^ through the definition of ***ε*** (13), where ***ξ***^*o*^ is the unique optimum in *Ω* (12) such that 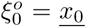. Recall that 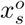 denotes the sensor concentration of ***ξ***^*o*^.

This proof involves finding the eigenvalues of the Jacobian and proving they are all negative. We will prove this by discussing the two distinct eigenspaces spaces that are qualitatively different that together span ℝ^*n−*1^. There is the (*n*−2)-dimensional eigenspace corresponding the 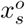-level-set. This entire space corresponds to eigenvalue −1. And there is the eigenspace that is transverse to this 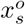-level-set, which has dimension 1 and can be shown to correspond to a negative eigenvalue.

The QSS 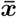 is a smooth function and a bijection on *E* (43) and the Jacobian matrix of 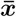 has maximal rank (Lemma 7). This implies that 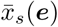 is a continuous function and the subset of *E* where 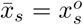 is a smooth manifold around ***ε***^*o*^. If we consider a small enough neighbourhood of the optimum ***ε***^*o*^, this manifold is approximated by an (*n* − 2)-dimensional linear space. Every element ***e*** in this space has 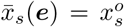, therefore its estimated optimum is correct,

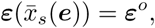

and it follows that that ***e*** is an eigenvector with eigenvalue −1: its time derivative is given by

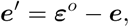

and in the linearisation around the optimum ***ε***^*o*^, the element ***e*** represents the vector ***e*** − *ε*^*o*^.

This accounts for *n* − 2 independent directions in the linearisation. Transverse to this there is either an eigenvector transverse to the *n* − 2-dimensional space, or a generalised eigenvector inside this same eigenspace. In the latter case all eigenvalues are −1 and the proof is complete. So we assume there is another eigenvector transverse to the space where 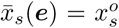. An element ***e*** that yields this eigenvector ***e** − **ε***^*o*^ therefore has 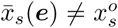. Assume without loss of generality that

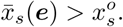

As ***e*** − ***ε***^*o*^ is an eigenvector, the direction of the time derivative

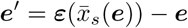

is found by multiplying it by its eigenvalue

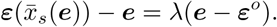

Hence the estimated enzyme distribution 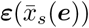 is on the line defined as containing ***e*** and ***ε***^*o*^

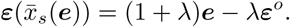

By Lemma 8, 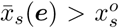 implies that 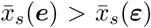, so the steady state level 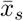 is decreased in the direction of the time derivative ***ε − e***. The steady state sensor level 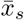 is increased in the direction of the vector time derivative vector ***e** − **ε***^*o*^. This opposing direction implies that the estimated ***ε*** is in the reverse direction on the line and therefore that *λ <* 0.

## 4 Global Stability of the shortest linear chain

The smallest linear chain that still has an internal metabolite is given by setting *n* = 2,

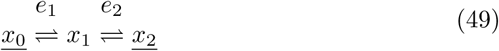

Since the QSS version of the *q*ORAC-controlled pathway in this case has only one dynamical variable, global stability is straightforward.

The differential algebraic system of our smallest model is given by

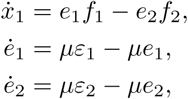

where the reaction kinetics are defined by the standard reversible MM kinetics,

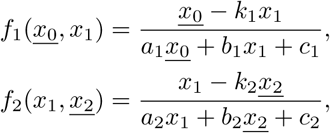

Furthermore,

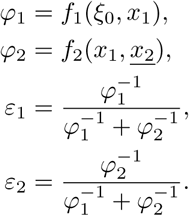

Here *x*_1_ is the concentration of the only internal metabolite and sensor for the nutrient concentration 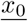. The estimated external concentration *ξ*_0_(*x*_1_) is the unique value such that *x*_1_ is the minimum of the objective function

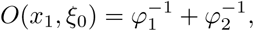

The nutrient concentration is positively ordered, so

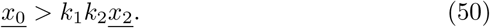

### 4.1 Timescale Separation

Note that the sum of enzyme *e*_1_ + *e*_2_ = 1 is conserved over time as

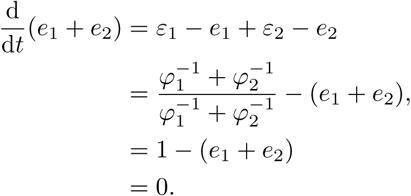

We eliminate *e*_2_ from the equations and set it to 1 − *e*_1_.

The nullcline 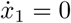 describes the slow manifold of the quasi-steady state, which according to Lemma 7 is a bijection between the interval 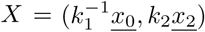 (44) and *E* = (*e*_1_, *e*_2_) : *e*_1_, *e*_2_ > 0, *e*_1_ + *e*_2_ = 1 (43).

In the smallest linear chain, the border of each of these two regions contains only two points and then the bijection can be extended to the border points as seen in the following lemma.

#### Lemma 10

*The nullcline* 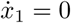 *starts at e*_1_ = 0, 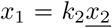 *and continues strictly increasing until e*_1_ = 1, 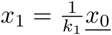.

*Proof* Considering 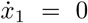, we get

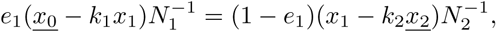

for *N*_1_, *N*_2_ the nonzero denominators of the flux functions *f*_1_ and *f*_2_ respectively.

If 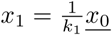, then 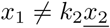 (50), so *e*_1_ = 1.

If 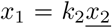 then 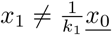 (50), so *e*_1_ = 0.

If 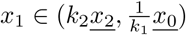, then *f*_1_, *f*_2_ > 0, hence the nullcline 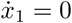 yields

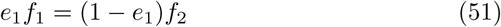

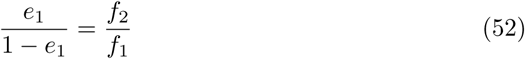

There is always a unique solution *e*_1_, as 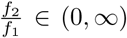 and 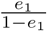 is increasing in *e*_1_, where (0, 1) is exactly mapped to (0, *∞*). The right hand side, 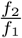 is increasing in *x*_1_ (*f*_2_ is increasing, *f*_1_ is decreasing), thus the unique solution for *e*_1_ increases as *x*_1_ increases.

#### Theorem 11

*The shortest linear chain with qORAC control is globally stable*.

*Proof* In the slow timescale, the internal metabolite *x*_1_ follows its nullcline 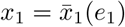 as given in Lemma 10.

The estimated optimal enzyme concentration

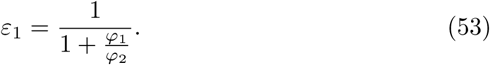

follows a similar expression of *x*_1_ as *e*_1_ does on the nullcline. Hence, 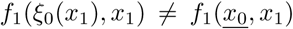 as in (52), but 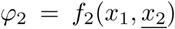. Now note that taking *ξ*_0_ instead of 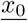 in *f*_1_ is the only difference between *e*_1_ on the slow manifold and *ε*_1_,

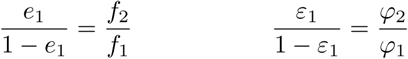

This leads us to the following three results:

– We can see the global uniqueness of the steady state solution. There is only one option for an element of the nullcline 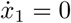 to also satisfy *e*_1_ = *ε*_1_. This is to have 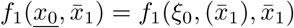. The function *f*_1_ is strictly increasing in its first argument, so the two sides can only be equal if 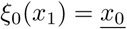, and as *ξ*_0_ is strictly increasing in *x*_1_, there is a unique solution.
– At its minimal value, 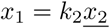, the estimate for the nutrient concentration is 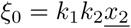, and then both flux functions are at equilibrium *f*_1_ = *f*_2_ = 0, leading to 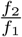 being undefined, but the limit exists and can be computed. For now, note that *e*_1_ has a limit value between 0 and 1. This limit is the optimal enzyme production continued all the way to equilibrium.
– At its maximum value, 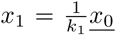, we have *ξ*_0_ at some value larger than 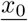 and it follows that unlike the slow manifold, *ε*_1_ > 0:

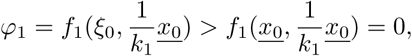

thus 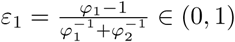

From the above remarks, we can conclude on the basis of continuity that the nullcline 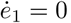, compared to the nullcline 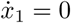, has higher *e*_1_ until the unique intersection, after which it has lower *e*_1_.

Hence, the dynamical system in the slow manifold

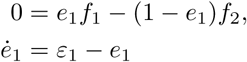

has a unique steady state and the flow is always directed towards this solution. Hence it is globally stable.

## 5 Discussion

In this paper we have studied the linear chain with reversible Michaelis-Menten kinetics, coupled to a set of equations for enzyme synthesis and dilution by growth which is designed to achieve maximal steady state flux. The construction is termed *q*ORAC. The design of this control does in no way guarantee that the coupled dynamical system is stable.

In (Planqué et al. 2018) we have given one example of a pathway coupled to *q*ORAC control in which dynamics do *not* converge to the optimal steady state but away from it. (This is the only counterexample known to date, and involves robustness to an internal kinetic parameter rather than robustness to external concentrations.)

The counterexample makes clear that *q*ORAC does not always work, and that an in-depth analysis is required to show at least local stability. In this paper we provide a proof of this for the simplest, but also arguable most important, metabolic pathway, the linear chain. To facilitate the analysis, we assumed that the chain is in QSS relative to enzyme synthesis, a reasonable biological assumption.

### 5.1 An outline of the construction

In the fast timescale, the metabolic network finds a globally unique and stable steady state ***x*** for fixed values of the enzyme concentrations ***e***. This metabolic steady state can be found for all possible enzyme concentration with total 1 (43), inducing the functions 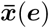 and 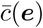. By implicit differentiation, we can find explicit partial derivatives of ***x*** to ***e*** (Corollary 5) and 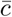 to ***e*** (Corollary 4).

The quasi-steady state 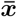 is shown to be a bijection between the set of conserved strictly positive enzyme concentrations *E* (43) and the set of positive metabolic concentrations where all flux functions are positive *X* (44). The inverse of this bijection is explicitly given in (45). Applying this to the expressions of the explicit partial derivatives for 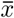 and 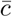 to ***e*** yields a dynamical system that is derived from the quasi-steady state assumption of the fast timescale and which only depends on the ***x*** variables; the dependency on ***e*** is eliminated.

Any estimate of the optimal steady state metabolic concentrations (i.e., an element of *Ω*) induces an optimal distribution of enzyme concentrations through the steady state equations of the metabolic network. Almost the same equations apply to this resultant enzyme distribution and its actual dual steady state metabolic concentrations. Comparing these two steady states yields the insight that the sensor concentration will change in the same direction as where the eventual optimum is found as we step from where we are now, to where the system is steering towards at this moment in time.

As a consequence, all but one eigenvalue of the Jacobi matrix are seen to be −1, and the local stability of the system becomes evident (Theorem 9).

### 5.2 Towards a Lyapunov Function

The argument that we used to prove global stability in the shortest chain relies on it becoming essentially a scalar problem. Therefore this proof cannot be extended to a linear chain of arbitrary length. However, it does show that the objective of showing global stability for the linear chain is potentially attainable. Thus we can try other methods that might extend to longer linear chains. So far, we have not been successful in this attempt. But we will introduce likely candidates for a Lyapunov function and show counterexamples.

Recall that a Lyapunov function has a negative time derivative everywhere except in the steady state, thereby showing that the steady state is globally stable.

Consider for a general linear chain the function

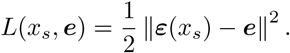

This shows some promise to be a Lyapunov function, because

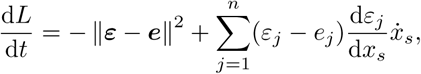

where a large part of the expression is obviously negative. If we can estimate the possibly positive part 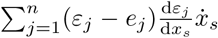 as less than the definitely negative part −∥***ε − e***∥^2^, then we can prove that *L* is a Lyapunov function.

However, we can find a counterexample. Through some judicious guessing, a set of parameters was chosen for the smallest linear chain (See Table 1). The Lyapunov function is the square of the Euclidian distance between the estimated optimal enzyme concentration vector ***ε*** and the current value ***e***. If we assume rapid equilibrium of the fast timescale, we will be on the slow manifold and move towards the unique steady state.

**Table 1:** Parameters that yield the counterexamples for the proposed Lyapunov functions. Note that because 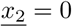, the parameters k_2_ and b_2_ are obsolete.

| Parameter | Figure 3 | Figure 4 |
| --- | --- | --- |
| $\underline{x}_0$ | 3 | 3 |
| $\underline{x}_2$ | 0 | 0 |
| $\underline{k}_1$ | 1 | 1 |
| $a_1$ | 0.8 | 5 |
| $b_1$ | 1.2 | 1.2 |
| $c_1$ | 0.05 | 0.05 |
| $a_2$ | 1.3 | 20 |
| $c_2$ | 6 | 6 |
| $\mu$ | 0.01 | 0.005 |

As can be immediately seen in Figure 3, this proposed Lyapunov function will then not necessarily be decreasing over time. If we start with very low *e*_1_, the rapid equilibrium will yield very low *x*_1_ and in the slow timescale, we will move upwards along the slow manifold, but the estimated enzyme concentration *ε*_1_ will move upward faster, increasing 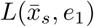 over time initially.

**Fig. 3:**
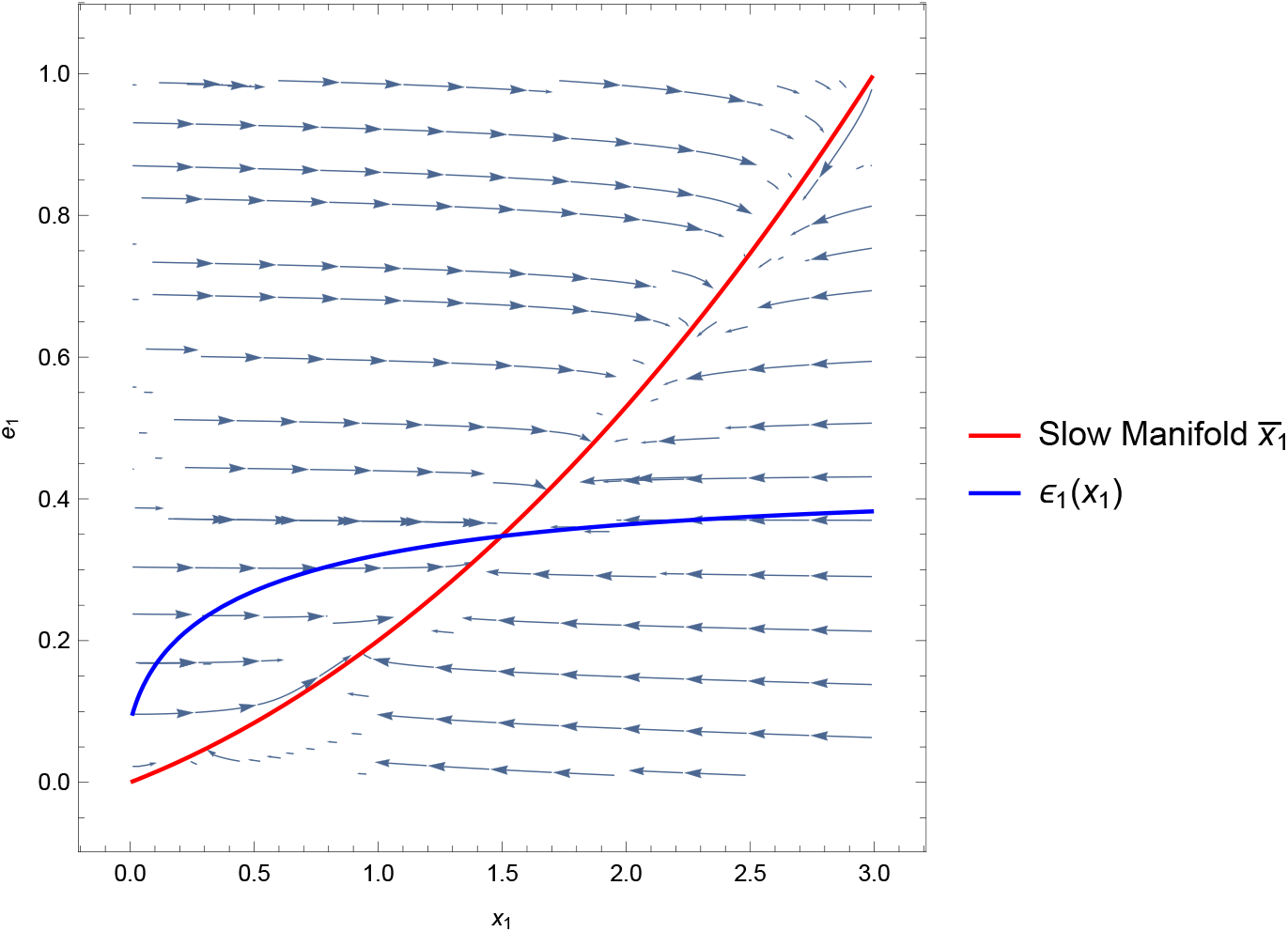
The dynamics of the smallest linear chain. The streamplot shows the actual dynamics for a small parameter µ. The red line describes the slow manifold and the blue line the estimated optimal enzyme concentration ε_1_ as a function of x_1_. This depicts a counterexample to the claim that ∥**ε − e**∥ can serve as a Lyapunov function.

**Fig. 4:**
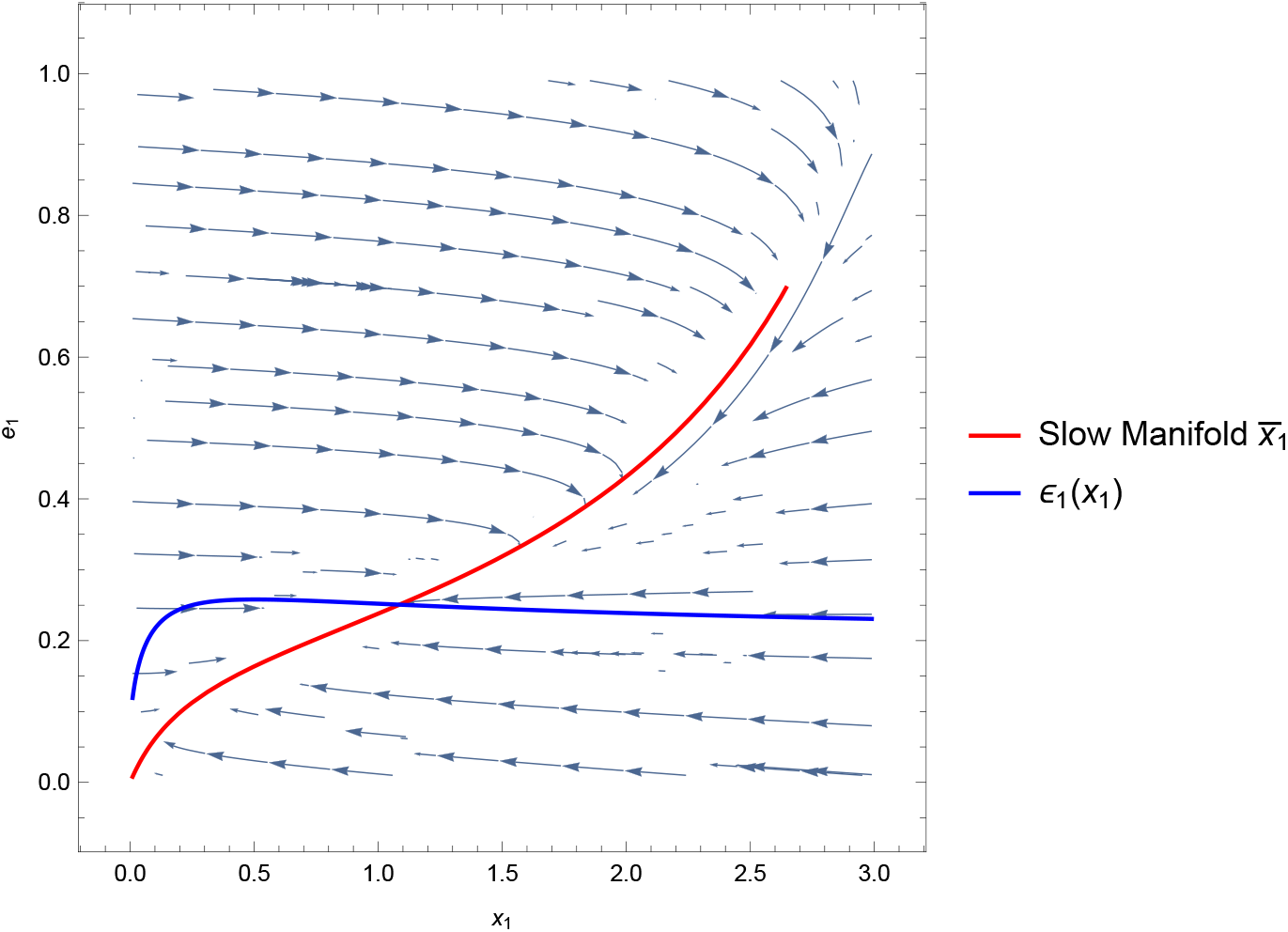
The dynamics of the smallest linear chain. The streamplot shows the actual dynamics for a small parameter µ. The red line describes the slow manifold and the blue line the estimated optimal enzyme concentration ε_1_ as a function of x_1_. This depicts a counterexample for that ∥**ε**^o^ − **ε**∥ can serve as a Lyapunov function.

We can also choose as second try,

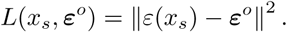

As the system flows in the slow timescale, this might seem like a good candidate for a Lyapunov function as the estimate will become better perhaps. Also for this choice, a counterexample can be found. Changing the parameters a bit from the previous counterexample (Table 1), the curve of *ε*_1_ shows a steeply increasing initial slope, followed by a very gently decreasing slope. As we follow 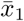 in the same manner as before, starting at *e*_1_ *≈* 0 (and thus *x*_1_ *≈* 0), *ε*_1_ will increase towards 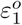 and then pass it, in order to gently decrease back to this eventual value. For a brief interval, this newly suggested candidate is increasing over time, rendering it of no use for a Lyapunov function.

A third candidate would be

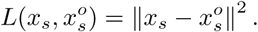

This is in fact a Lyapunov function for the smallest linear chain, but in a linear chain with 3 enzymes, we already found a counterexample (not illustrated).

The only remaining candidate for a Lyapunov function we have considered for which we have found no counterexamples to date is

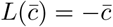

minus the steady state flux as the system flows in the slow manifold (or equivalently the objective function). In every numerical simulation that we have performed, the steady state flux 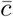 would improve over time in a monotone manner. Proving that this is indeed a Lyapunov function is another matter. By taking a great deal of explicit information about the control and the quasisteady state into account, in the Appendix (Theorem 12) we show that

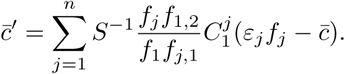

We have not been able to take this result further, and show it to have a definite sign.

## 6 Appendix

## 6.1 Inverse Quasi Steady State; Enzyme levels follow the metabolic concentrations

Lemma 7 shows that 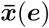 is a smooth bijection and its inverse *ē* is explicitly defined by (45). Furthermore, we can use these explicit steady states ***e*** to derive expressions for 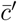 and 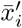, *i* = 1, …, *n* − 1, that only depend on ***x***, such that we have simplified the total dynamical system to having only the metabolic concentrations ***x*** as variables. This is based on the resulting expressions for 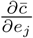 from Lemma 4 and 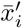 from Corollary 6. Note that from the definition of the system in Section 3.1.2 it is not obvious that such a construction is possible, but now that we have constructed the derivatives of 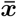 and 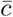 and have equations that express ***e*** in terms of ***x***, it follows immediately.

**Theorem 12** *The steady state flux has time derivative in the slow timescale*

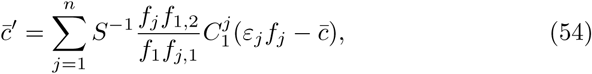

*where*

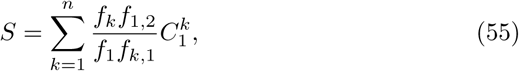

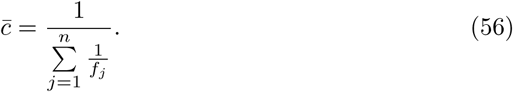

*Proof* Using the expression for 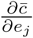 (31), we see

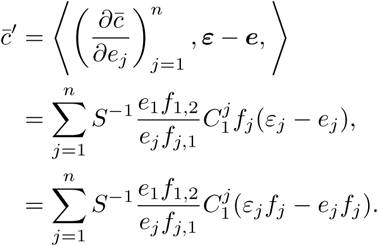

An alternative form of the steady state equations

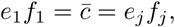

can be written as

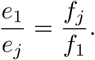

Substituting these forms into the expression for 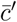 almost eliminates the enzyme concentrations from the equation,

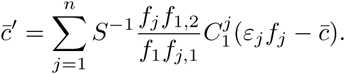

The only proliferance of the *e_j_* is in 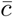 and

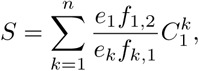

so through the steady state equations this can be rewritten without the enzyme concentrations,

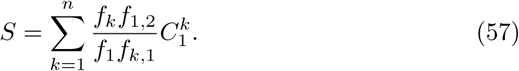

Furthermore we can write 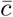 as a function of only ***x***, because

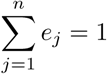

and from the steady state equations

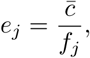

so we can see

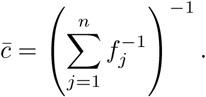

The time dependence of the metabolic concentrations ***x**^′^* in terms of only ***x*** follows by taking Corollary 6, while substituting any instance of *e*_*j*_ by using the quasi-steady state equations (45).

### Theorem 13

*By means of the quasi-steady state assumption, the system can be rewritten as an ODE system in only* 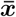 *by*

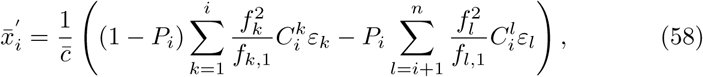

*where P*_*i*_ *is denoted in equation* (59).

*Proof* Start with the following substitution from the quasi-steady state:

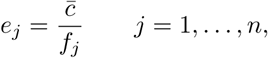

where 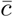 is given in (56)

We then get an expression of the time dependence of ***x*** solely in terms of

***x***,

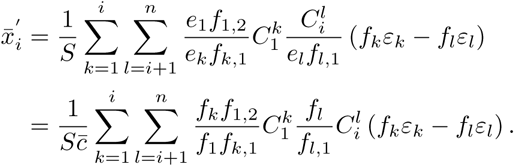

Recall that *S* can be rewritten through the steady state equations to not depend on ***e*** (57).

We can rewrite the expression for 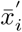, substituting 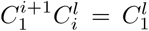 on the second line,

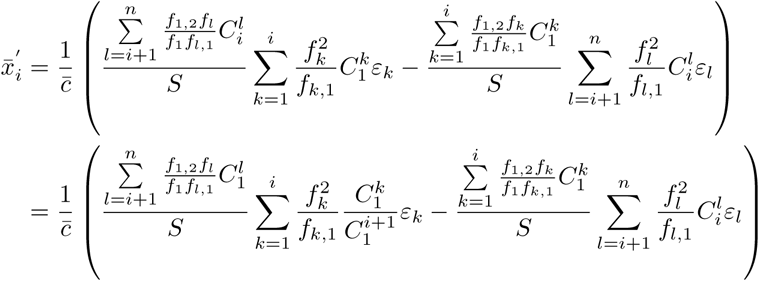

Above we recognise the two complementary parts of the total sum of *S* divided by *S* (55). So the following is always a fraction between 0 and 1 *P*_*i*_ ∈ [0, 1],

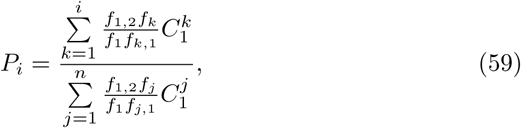

which can be substituted in the expression,

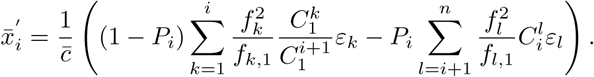

Recall that we introduced a natural extention to the definition of 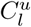 in (20). From this we can see, if *k ≤ i*,

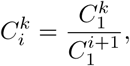

which leads to the expression

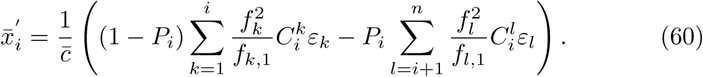

## 6.2 Global stability of the linear chain with fixed enzyme concentrations

Here we provide a new proof for the global stability of the linear chain with fixed enzyme concentrations,

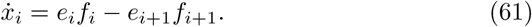

For the purposes of this Appendix, the Michaelis-Menten kinetics are chosen here to have the following form,

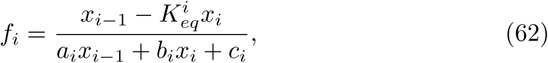

An older proof, which seems to have been largely forgotten, can be found in (Smillie 1984).

### Theorem 14

*The unique steady state of (61)–(62) is globally stable*.

*Proof* Write the system in the form

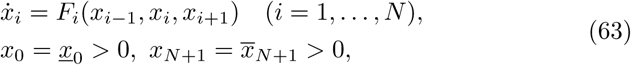

where

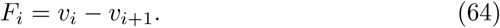

Note that

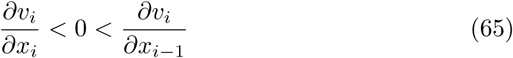

for all nonnegative values of the concentrations.

The proof follows from sub- and supersolution arguments and a suitable comparison principle. We show below that (65) implies that two solutions with initial data satisfying 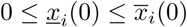 for *i* = 1, …,*N* have the property that 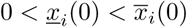 for all *t >* 0 and all *i* = 1, …, *n*, unless 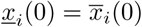 for all *i* = 1, …, *N*. (The reader should not be confused by the different use of the underlines in the notation.) We use this comparison principle with small and large initial data, both characterised as being a thermodynamic equilibrium for the linear chain from *x*_1_ to *x*_*N*_, defined by

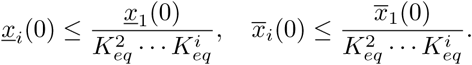

We can then take 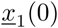 so close to zero and 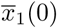 so large that

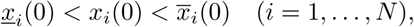

whence for all *t >* 0

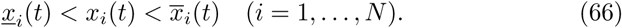

These solutions are called the subsolution, the solution and the supersolution. We then also show that the sub- and the super solution satisfy

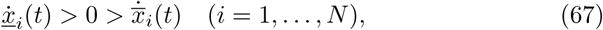

for all *t >* 0 whence both of them, and therefore also the solution itself converge to the unique steady state determined by 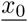 and 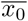. Below we give some of the details of the argument.

Writing 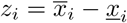 we have

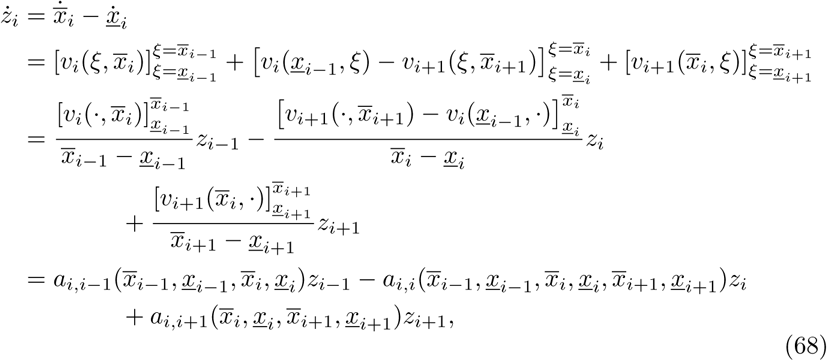

with the obvious adaptation for *i* = 1 and *i* = *N*. Note that via long division the coefficients *a*_*i,i* − 1_, *a*_*i,i*_, *a*_*i,i*+1_ of *z*_*i* − 1_, *z*_*i*_, *z*_*i*+1_ are smooth positive rational functions of their arguments, because of (64)–(65). The strict comparison principle now follows by standard arguments: if the initial data are strictly ordered, then at a first *t* for which there exists *i* with *z*_*i*_ = 0 we must have that the neighbouring *z*_*j*_ are also zero and hence 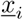 and 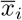 coincide. Repeating the argument it follows that all *z*_*j*_ are zero at this value of *t*. Thus 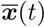 and 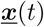 coincide, contradicting the fundamental theorem about existence and uniqueness for solutions of initial value problems systems of ODE’s. Thus such a first *t* cannot exist.

Solutions with initial data 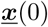, 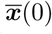 which are ordered, but not strictly ordered in the sense that some of but not all the initial concentrations coincide, can be approximated with solutions with strictly ordered initial data 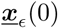, 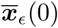. Taking the limit it follows that the *z*_*i*_ are globally nonnegative. But then (68) implies 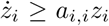 and the real analyticity of solutions implies that each *z*_*i*_(*t*) is either globally positive, or identically equal to zero. Any such zero solution can only have zero neighbours because otherwise, again in view of (68), its derivative would be positive. Thus all *z*_*i*_(*t*) are strictly positive because at least must be, as the fundamental theorem prohibits all *z*_*i*_(*t*) to vanish.

It remains to be shown that all 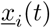 and 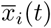 are strictly monotone in the sense of (67). From 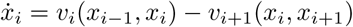 we obtain

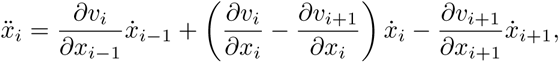

i.e.,

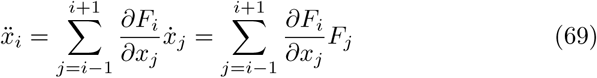

with

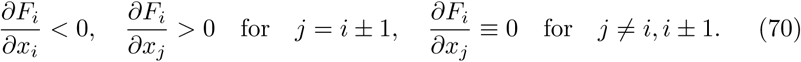

Suppose that initially *F*_*i*_(***x***) *<* 0 for all *i* = 1, …, *N* and for some first *t* there is an *i* with 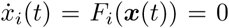. Then by (69) and (70) we have 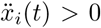 unless both *F*_*i*_−_1_(***x***(*t*)) and *F*_*i*+1_(***x***(*t*)) also vanish. If *i* = 1 this applies only to *F*_2_ and if *i* = *N* only to *F*_*N*+1_ of course. But then it follows by the same argument that all *F*_*i*_(***x***(*t*)) vanish, contradicting the fundamental theorem. The same argument applies to solutions for which initially all *F*_*i*_(***x***) > 0.

The strong version of this invariance, assuming only *F*_*i*_(***x***) ≤ 0 for all *i* = 1, …, *N* with at least one inequality strict, seems a bit trickier to establish, so we cannot directly conclude that the derivatives of 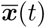 and 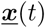 are strictly negative and strictly positive. Using a variation of the above argument for existence of a unique steady state, the initial data for both these solutions can easily be perturbed to initial data for which 0 = *v*_1_ *< … < v*_*N*_ = *ϵ*, for which *ϵ* = *v*_2_ *> … > v*_*N*+1_ = 0. The corresponding solutions 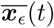 and 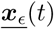 are then globally strictly ordered and strictly monotone in the sense that the derivatives are nonzero. Clearly we can take the limit *ϵ* → 0 and conclude that 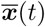 and 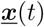 are then globally strictly ordered and monotone in the sense that the derivatives do not change sign. But then we also have that for each *i*

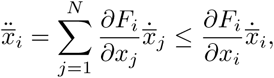

whence 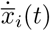 has the property that once it is strictly negative it remains strictly negative. But since all 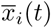 are real analytic in *t* we have that either 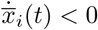 for all *t >* 0 or 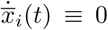. Hence, the latter case is excluded by the same argument as used above to establish the global strict positivity of all 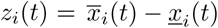.

## References

M. Basan, S. Hui, H. Okano, Z. Zhang, Y. Shen, J. R. Wiliamson, and T. Hwa. Overflow metabolism in E. coli results from efficient proteome allocation. Nature, 528:99–104, 2015.

E. Bosdriesz, D Molenaar, B. Teusink, and F. J. Bruggeman. How fast-growing bacteria robustly tune their ribosome concentration to approximate growth-rate maximisation. FEBS Journal, 282(10):2029–44, 2015.

A. Cornish-Bowden. Fundamentals of Enzyme Kinetics. Wiley-Blackwell, 4th edition, 2004.

E. Dekel and U. Alon. Optimality and evolutionary tuning of the expression level of a protein. Nature, 436:588–592, 2005.

W. Gilbert and B. Müller-Hill. Isolation of the lac repressor. Proc. Nat. Acad. Sciences USA, 56(6):1891–1898, 1966.

P. R. Jensen, O. Michelsen, and H. V. Westerhoff. Experimental determination of control by the H^+^-ATPase in Escherichia coli. J. Bioenerg. Biomem., 27: 543–554, 1995.

L. Keren, J. Hausser, M. Lotan-Pompan, I. Vainberg Slutskin, H. Alisar, S. Kaminski, A. Weinberger, U. Alon, R. Milo, and E. Segal. Massively parallel interrogation of the effects of gene expression fevels on fitness. Cell, 166(5):1282–1294.e18, 2016.

Karl Kochanowski, Benjamin Volkmer, Luca Gerosa, Bart R Haverkorn van Rijsewijk, Alexander Schmidt, and Matthias Heinemann. Functioning of a metabolic flux sensor in Escherichia coli. Proc. Nat. Acad. Sciences USA, 110(3):1130–1135, January 2013.

O. Kotte, J. B. Zaugg, and M. Heinemann. Bacterial adaptation through distributed sensing of metabolic fluxes. Mol. Syst. Biol., 6:355, 2010.

O Kotte, B Volkmer, J L Radzikowski, and M Heinemann. Phenotypic bistability in Escherichia coli’s central carbon metabolism. Mol. Syst. Biol., 10 (7):736–736, July 2014.

G. W. Li, D. Burkhardt, C. Gross, and J. S. Weissman. Quantifying absolute protein synthesis rates reveals principles underlying allocation of cellular resources. Cell, 157(3):624–635, 2014.

S. Müller, G. Regensburger, and R. Steuer. Enzyme allocation problems in kinetic metabolic networks: Optimal solutions are elementary flux modes. J. Theor. Biolog, 347:182–190, 2014.

E. Noor, A. Flamholz, A. Bar-Even, D. Davidi, R. Milo, and W. Liebermeister. The protein cost of metabolic fluxes: Prediction from enzymatic rate laws and cost minimization. PLoS Comp. Biol., 12(11):e1005167, 2016.

J. A. Papin, J. Stelling, N. D. Price, S. Klamt, S. Schuster, and B. Ø. Palsson. Comparison of network-based pathway analysis methods. Trends in Biotechnology, 22:400–405, 2004.

R. Planqué, J. Hulshof, B. Teusink, J. C. Hendriks, and F. J. Bruggeman. Maintaining maximal metabolic flux by gene expression control. PLoS Comp. Biol., 14(9):e1006412, 2018. https://doi.org/10.1371/journal.pcbi.1006412.

M. Schaechter, J. L. Ingraham, and F. C. Neidhardt. Microbe. ASM Press, Washington DC, 2006.

S. Schuster and C. Hilgetag. On elementary flux modes in biochemical reaction systems at steady state. J. Biol. System, 2:165–185, 1994.

S. Schuster, C. Hilgetag, J. H. Woods, and D. A. Fell. Reaction routes in bio-chemical reaction systems: algebraic properties, validated calculation procedure and example from nucleotide metabolism. J. Math. Biolog, 45: 153–181, 2002.

M. Scott, S. Klumpp, E. M. Mateescu, and T. Hwa. Emergence of robust growth laws from optimal regulation of ribosome synthesis. Mol. Syst. Biol., 10:747, 2014.

C. A. Sellick, R. N. Campbell, and R. J. Reece. Galactose metabolism in yeast—structure and regulation of the leloir pathway enzymes and the genes encoding them. Int. Rev. Cell. Mol. Biol., 269:111–150, 2008.

J. Smillie. Competitive and cooperative tridiagonal systems of differential equations. SIAM J. Math. Anal., 15(3):530–534, 1984.

M. T. Wortel, H. Peters, J. Hulshof, B. Teusink, and F. J. Bruggeman. Metabolic states with maximal specific rate carry flux through an elementary flux mode. FEBS Journal, 281:1547–1555, 2014.

C. You, H. Okano, S. Hui, Z. Zhang, M. Kim, C. W. Gunderson, Y.-P. Wang, P. Lenz, D. Yan, and T. Hwa. Coordination of bacterial proteome with metabolism by cyclic AMP signalling. Nature, 500:301–306, 2013.

